## Supplementary Figure Legends for "Genetic Inactivation of the β1 adrenergic receptor prevents Cerebral Cavernous Malformations in zebrafish"

Supplementary Figure 1. (A) *adrb1^-/-^* embryos displayed a decrease of heart rate compared to wild type embryos at 28hpf. Unpaired two-tailed t test was performed and P<0.0001. (B) *adrb1^-/-^* embryos showed significant decrease of RBC velocity compared to wild type embryos. Time-lapses on a single z-plane was performed at the frequency of 160.97ms/frame (372 frames/minute) on Fast Airyscanning. 10 embryos from each group were scanned, and 3 red blood cells were traced from each embryo. The measurement was performed using ImageJ. Unpaired two-tailed t-test, P=0.0346.

Supplementary Figure 2. 2,3-BDM decreases the heart rate in 30hpf zebrafish embryos. Two-tailed paired t-test was used for statistical analysis. P<0.0001.

Supplementary Figure 3. No significant difference of CVP development was observed between *adrb1^-/-^* and wild type embryos. (A and B) Representative bright field pictures of *adrb1^-/-^* (A) and wild type (B) embryos at 36hpf. Scale bar: 500 µm. (C and D) Representative confocal pictures of CVP in *adrb1^-/-^* (C) and wild type (D) embryos at 36hpf. Red and blue brackets indicate the aorta and CVP respectively. Scale bar:100µm.

Supplementary Video 1. The cardiac pumping is perturbed in *adrb1^-/-^* embryos at 28hpf. *adrb1^-/-^* embryos displayed weaker heartbeat compared to wild type embryos at 28hpf.

Supplementary Video 2. The blood flow in CVP is perturbed in *adrb1^-/-^* embryos compared to wild type embryos at 28hpf.

Supplementary Table 1. Comparison of the two-phase zebrafish CCM model with mouse CCM model and human CCM. “?” means it is yet to be determined. “-” means it is not applicable. “1”. Perilesional red blood cell leakage was seen.

Supplementary Table 2. The predicted off-targets genomic sites produced by *adrb1* CRISPR. These genomic sites were sequenced and found no mutations. Primer sequence used for amplifying these sites were listed.
