## Supplementary Figures and Tables for "Genetic Inactivation of the β1 adrenergic receptor prevents Cerebral Cavernous Malformations in zebrafish"

### Slide 1
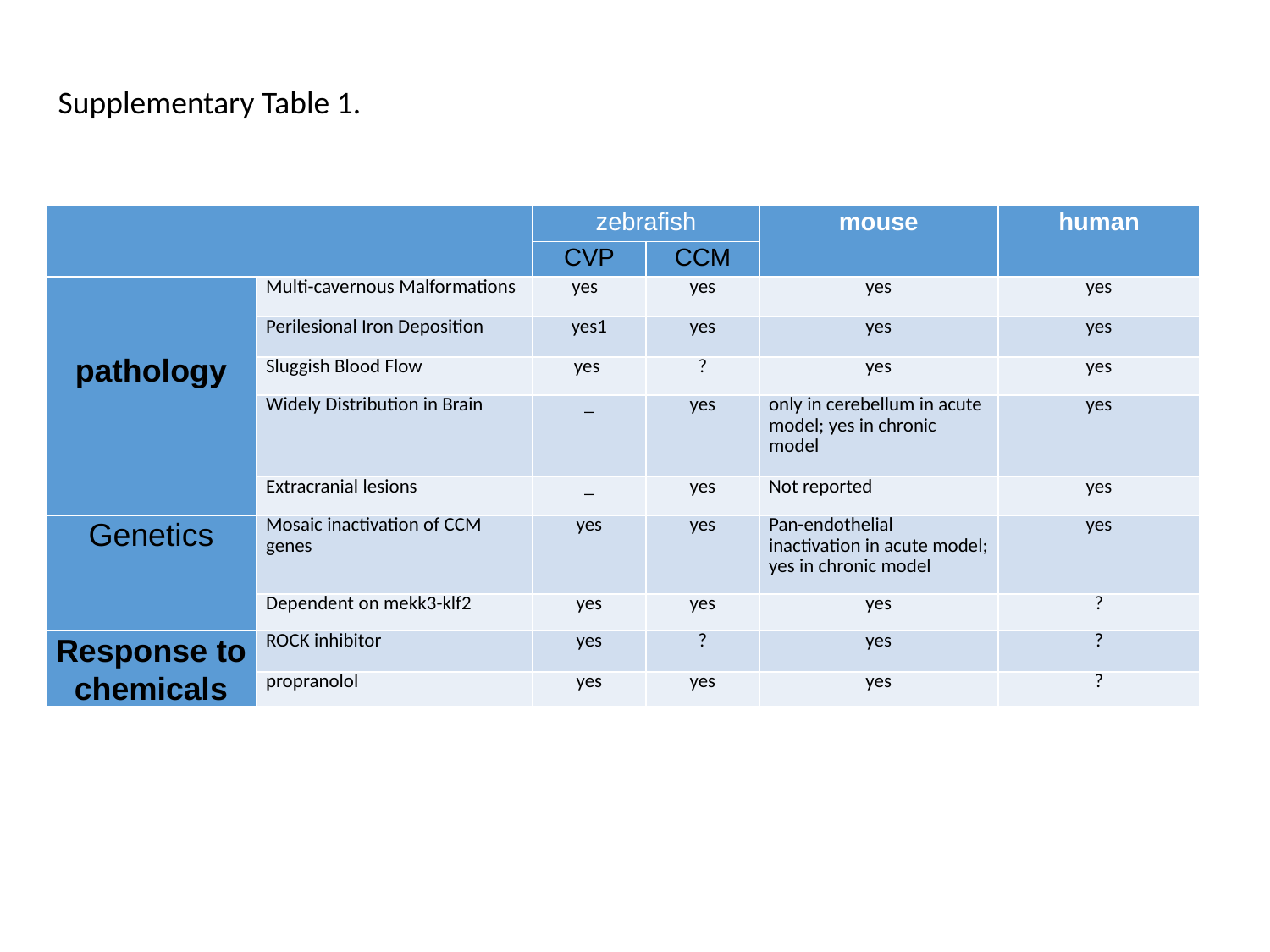

Supplementary Table 1.
| | | zebrafish | | mouse | human |
| --- | --- | --- | --- | --- | --- |
| | | CVP | CCM | | |
| pathology | Multi-cavernous Malformations | yes | yes | yes | yes |
| | Perilesional Iron Deposition | yes1 | yes | yes | yes |
| | Sluggish Blood Flow | yes | ? | yes | yes |
| | Widely Distribution in Brain | \_ | yes | only in cerebellum in acute model; yes in chronic model | yes |
| | Extracranial lesions | \_ | yes | Not reported | yes |
| Genetics | Mosaic inactivation of CCM genes | yes | yes | Pan-endothelial inactivation in acute model; yes in chronic model | yes |
| | Dependent on mekk3-klf2 | yes | yes | yes | ? |
| Response to chemicals | ROCK inhibitor | yes | ? | yes | ? |
| | propranolol | yes | yes | yes | ? |

### Slide 2
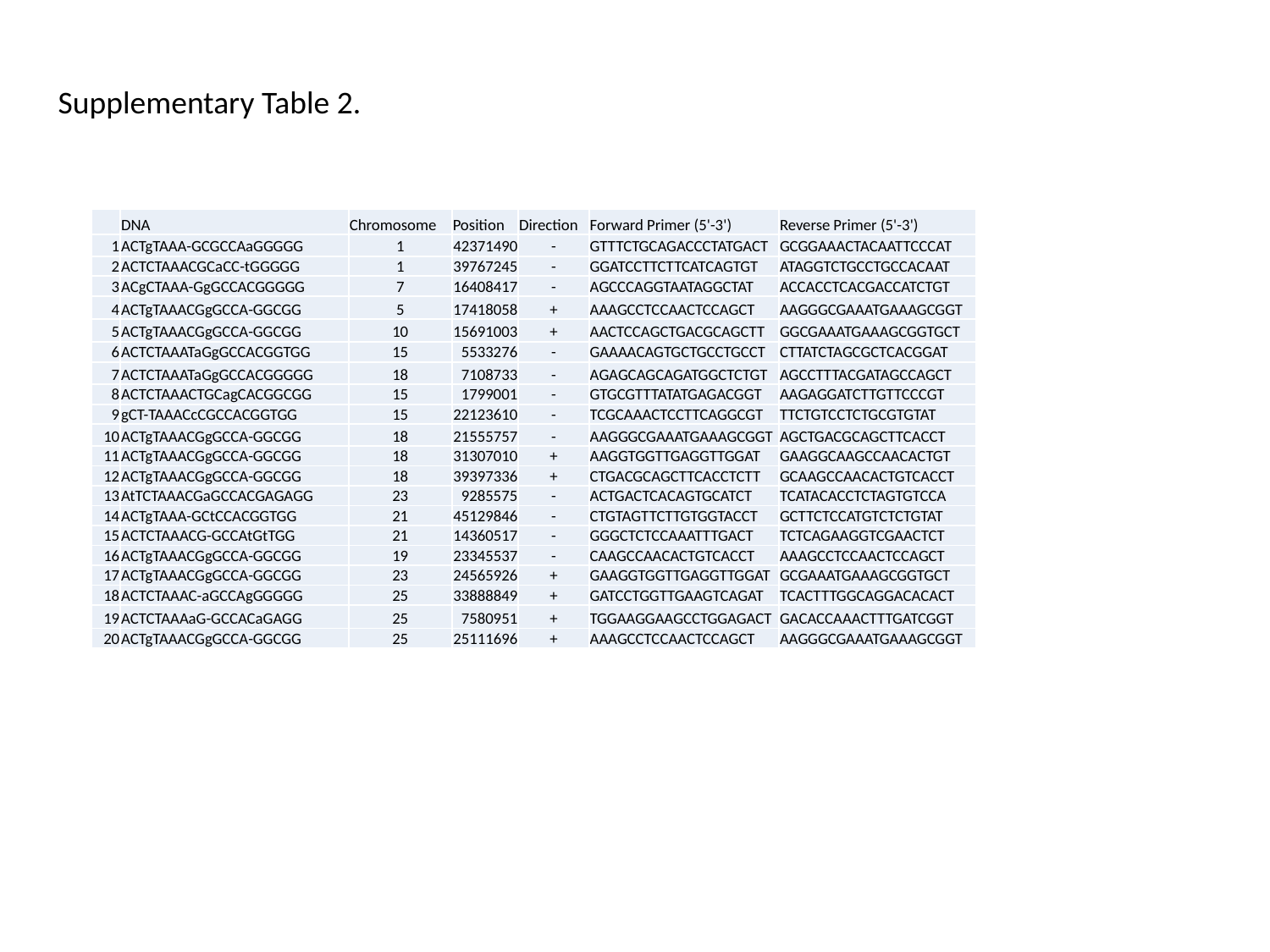

Supplementary Table 2.
| | DNA | Chromosome | Position | Direction | Forward Primer (5'-3') | Reverse Primer (5'-3') |
| --- | --- | --- | --- | --- | --- | --- |
| 1 | ACTgTAAA-GCGCCAaGGGGG | 1 | 42371490 | - | GTTTCTGCAGACCCTATGACT | GCGGAAACTACAATTCCCAT |
| 2 | ACTCTAAACGCaCC-tGGGGG | 1 | 39767245 | - | GGATCCTTCTTCATCAGTGT | ATAGGTCTGCCTGCCACAAT |
| 3 | ACgCTAAA-GgGCCACGGGGG | 7 | 16408417 | - | AGCCCAGGTAATAGGCTAT | ACCACCTCACGACCATCTGT |
| 4 | ACTgTAAACGgGCCA-GGCGG | 5 | 17418058 | + | AAAGCCTCCAACTCCAGCT | AAGGGCGAAATGAAAGCGGT |
| 5 | ACTgTAAACGgGCCA-GGCGG | 10 | 15691003 | + | AACTCCAGCTGACGCAGCTT | GGCGAAATGAAAGCGGTGCT |
| 6 | ACTCTAAATaGgGCCACGGTGG | 15 | 5533276 | - | GAAAACAGTGCTGCCTGCCT | CTTATCTAGCGCTCACGGAT |
| 7 | ACTCTAAATaGgGCCACGGGGG | 18 | 7108733 | - | AGAGCAGCAGATGGCTCTGT | AGCCTTTACGATAGCCAGCT |
| 8 | ACTCTAAACTGCagCACGGCGG | 15 | 1799001 | - | GTGCGTTTATATGAGACGGT | AAGAGGATCTTGTTCCCGT |
| 9 | gCT-TAAACcCGCCACGGTGG | 15 | 22123610 | - | TCGCAAACTCCTTCAGGCGT | TTCTGTCCTCTGCGTGTAT |
| 10 | ACTgTAAACGgGCCA-GGCGG | 18 | 21555757 | - | AAGGGCGAAATGAAAGCGGT | AGCTGACGCAGCTTCACCT |
| 11 | ACTgTAAACGgGCCA-GGCGG | 18 | 31307010 | + | AAGGTGGTTGAGGTTGGAT | GAAGGCAAGCCAACACTGT |
| 12 | ACTgTAAACGgGCCA-GGCGG | 18 | 39397336 | + | CTGACGCAGCTTCACCTCTT | GCAAGCCAACACTGTCACCT |
| 13 | AtTCTAAACGaGCCACGAGAGG | 23 | 9285575 | - | ACTGACTCACAGTGCATCT | TCATACACCTCTAGTGTCCA |
| 14 | ACTgTAAA-GCtCCACGGTGG | 21 | 45129846 | - | CTGTAGTTCTTGTGGTACCT | GCTTCTCCATGTCTCTGTAT |
| 15 | ACTCTAAACG-GCCAtGtTGG | 21 | 14360517 | - | GGGCTCTCCAAATTTGACT | TCTCAGAAGGTCGAACTCT |
| 16 | ACTgTAAACGgGCCA-GGCGG | 19 | 23345537 | - | CAAGCCAACACTGTCACCT | AAAGCCTCCAACTCCAGCT |
| 17 | ACTgTAAACGgGCCA-GGCGG | 23 | 24565926 | + | GAAGGTGGTTGAGGTTGGAT | GCGAAATGAAAGCGGTGCT |
| 18 | ACTCTAAAC-aGCCAgGGGGG | 25 | 33888849 | + | GATCCTGGTTGAAGTCAGAT | TCACTTTGGCAGGACACACT |
| 19 | ACTCTAAAaG-GCCACaGAGG | 25 | 7580951 | + | TGGAAGGAAGCCTGGAGACT | GACACCAAACTTTGATCGGT |
| 20 | ACTgTAAACGgGCCA-GGCGG | 25 | 25111696 | + | AAAGCCTCCAACTCCAGCT | AAGGGCGAAATGAAAGCGGT |

### Slide 3
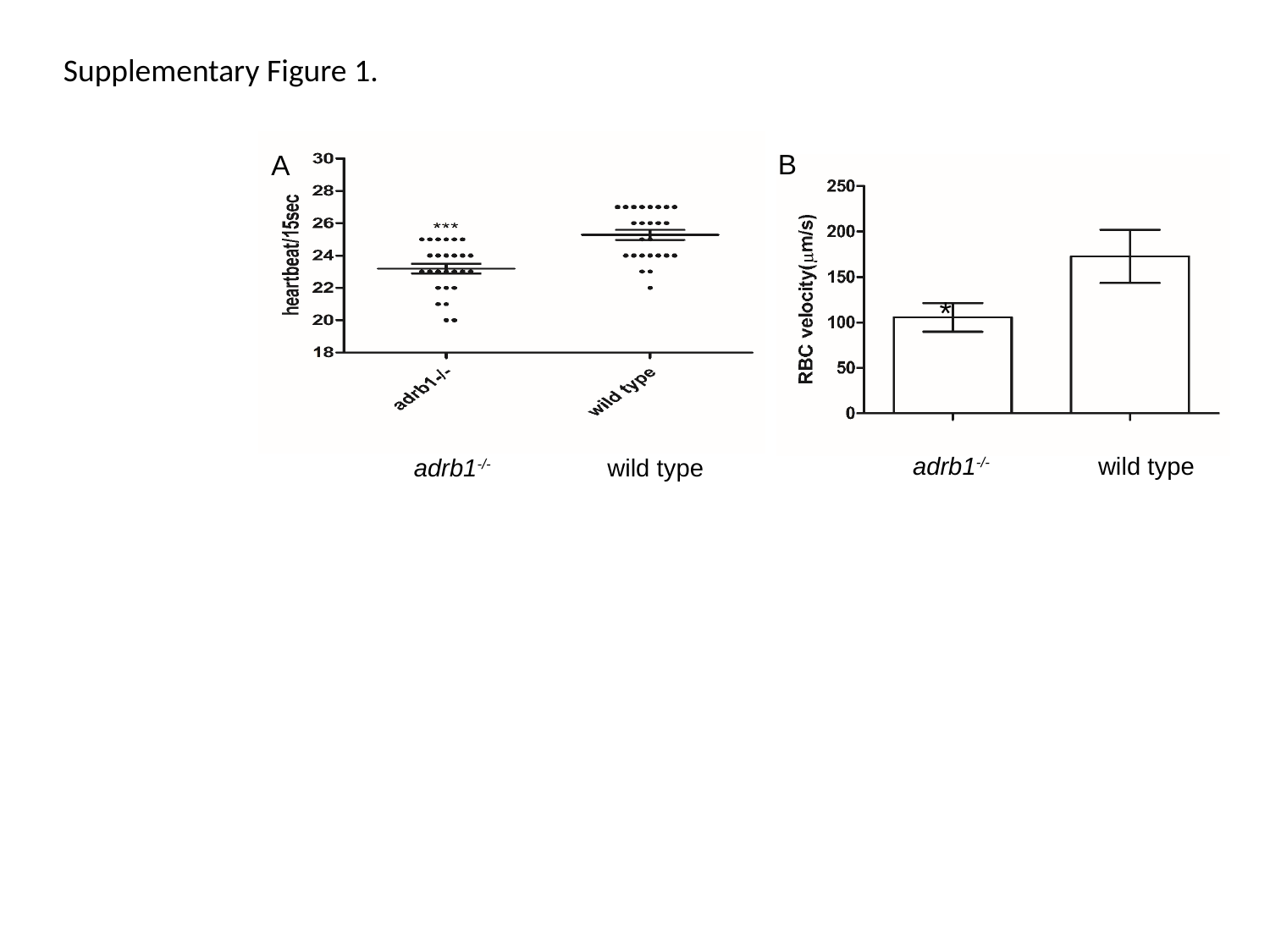

Supplementary Figure 1.
B
A
*
adrb1^-/-^
wild type
adrb1^-/-^
wild type

### Slide 4
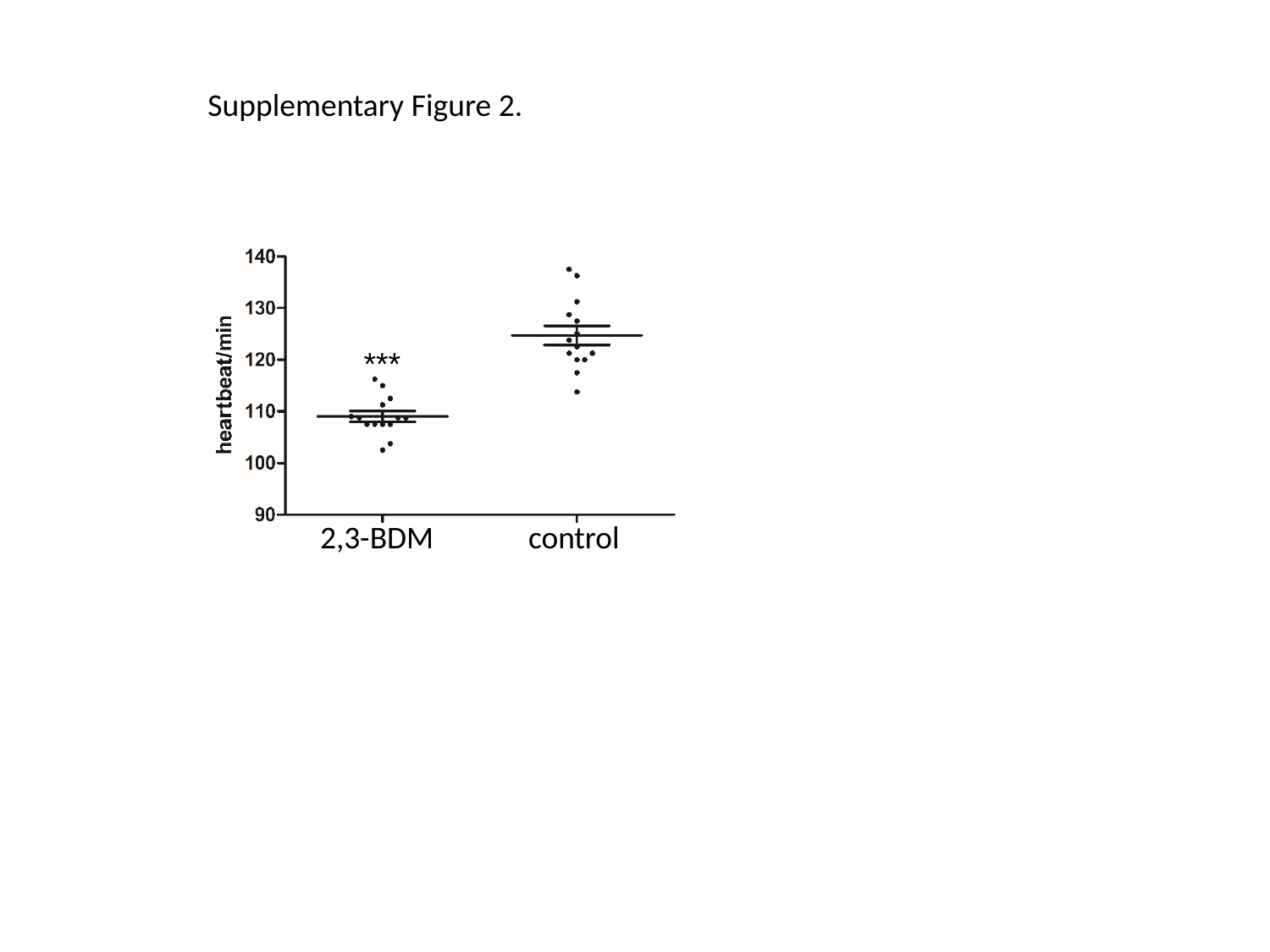

Supplementary Figure 2.
***
2,3-BDM
control

### Slide 5
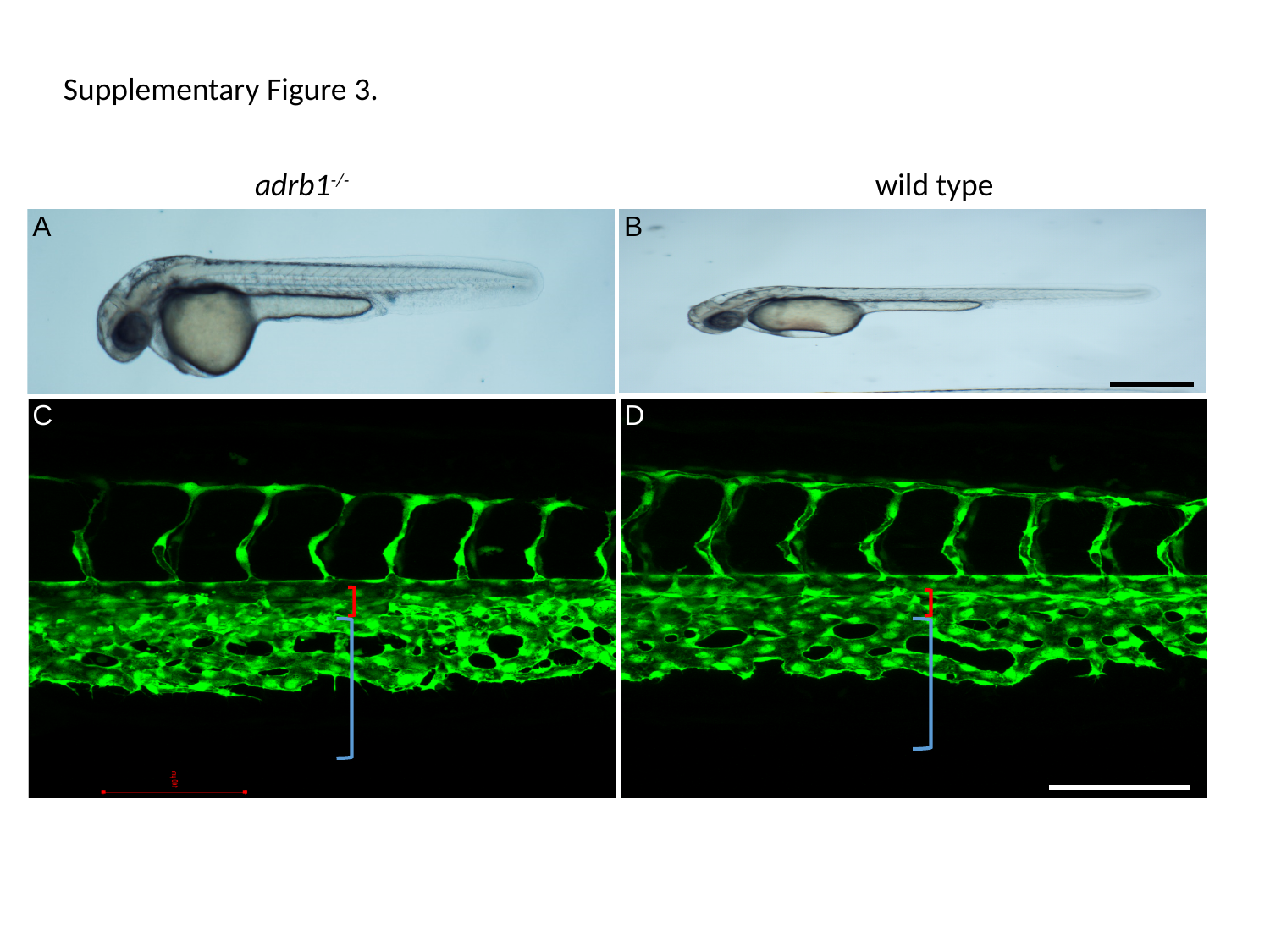

Supplementary Figure 3.
adrb1^-/-^
wild type
A
B
C
D
